# Refinement of AlphaFold-Multimer structures with single sequence input

**DOI:** 10.1101/2022.12.27.521991

**Authors:** Toshiyuki Oda

## Abstract

AlphaFold2, introduced by DeepMind in CASP14, demonstrated outstanding performance in predicting protein monomer structures. It could model more than 90% of targets with high accuracy, and so the next step would surely be multimer predictions, since many proteins do not act by themselves but with their binding partners. After the publication of AlphaFold2, DeepMind published AlphaFold-Multimer, which showed excellent performance in predicting multimeric structures. However, its accuracy still has room for improvement compared to that of monomer predictions by AlphaFold2. In this paper, we introduce a fine-tuned version of AlphaFold-Multimer, named AFM-Refine-G, which uses structures predicted by AlphaFold-Multimer as inputs and produces refined structures without the help of multiple sequence alignments or templates. The performance of AFM-Refine-G was assessed using four datasets: Ghani_et_al_Benchmark2 and Yin_et_al_Hard using AlphaFold-Multimer version 2.2 outputs, and CASP15_multimer and Yin_and_Pierce_af23 using AlphaFold-Multimer version 2.3 outputs. Of 1925 predicted structures, 203 had DockQ improvement > 0.05 after refinement, demonstrating that our model is useful for the refinement of multimer structures. However, considering the per target success rate, the overall improvement was modest, suggesting that the original AlphaFold-Multimer network had already learned a biophysical energy function independent of MSAs or templates, as proposed by Roney and Ovchinnikov (Roney and Ovchinnikov, 2022). Furthermore, both the default AlphaFold-Multimer and our refinement model showed lower performance for immune-related targets compared to general targets, indicating that room for improvement remains.

**Availability:** The inference scripts are available from https://github.com/t-oda-ic/afm_refiner under the Apache License, Version 2.0. The network parameters are available from https://figshare.com/articles/online_resource/afm_refine_g_20230110_zip/21856407 under the license CC BY 4.0.

## Introduction

Predicting the protein structure has long been a major challenge in biology. AlphaFold2, presented by DeepMind in CASP14, demonstrated outstanding performance, and could model > 90% (GDT_TS > 70 for 87 out of 92 domains) of the CASP14 targets with high accuracy (Jumper, et al., 2021; Jumper, et al., 2021; Pereira, et al., 2021). After the competition, its derivative: AlphaFold-Multimer, was published (Evans, et al., 2022), and also showed excellent performance in multimer prediction (Evans, et al., 2022; Yin, et al., 2022).

However, unlike the performance of AlphaFold2 when predicting monomer structure, room for improvement remains for multimer predictions. The success (including not only ‘High’ but also lower quality classes in CAPRI (Lensink, et al., 2007) criteria) rate is approximately 85% for general heterodimer complexes and 25% for immune-related complexes or repebody-antigen complexes (Yin, et al., 2022). Predicting the multimeric structure of proteins is as important as that of the monomeric structure because many proteins do not act alone and require their binding partners.

We assumed that there are two types of problems associated with the difficulty in predicting multimer structures: 1) Many modern protein structure prediction methods (which also seems to be true for AlphaFold2 and AlphaFold-Multimer) rely on co-evolution information between residues (Anishchenko, et al., 2021; Zheng, et al., 2021). However, to extract co-evolution information between residues in different proteins, it is necessary to find the correct sequence pairs in multiple sequence alignment (MSA). The pairing is easy when no paralogs are present or if other information (e.g., they are coded in the same operon) can be used to identify the interacting partner. Unfortunately, pairing is usually difficult because many proteins have paralogs and the number of known operons is limited. To tackle this problem, several techniques have been used (Feinauer, et al., 2016; Humphreys, et al., 2021; Ovchinnikov, et al., 2014) but the procedure cannot be expected to be perfect, and inaccurate pairings add noise to the prediction. (We should note that, at this point, although the number of cases is limited, Gao et al. showed that AlphaFold-Multimer without pairing improved the rate of prediction success (Gao, et al., 2022).) 2) Amino acid profiles constructed from evolutionarily related sequences in MSAs are useful for structure prediction (Rost and Sander, 1995). This is because evolutionarily related sequences, often referred to as family or superfamily, have similar structures in most cases (Chandonia, et al., 2022; Sillitoe, et al., 2021). Therefore, hydrophilic residues found in the same regions of evolutionarily related sequences would support the prediction that these regions are solvent accessible, and vice versa. However, sequence-specific regions, such as Complementarity Determining Regions (CDRs) of antibodies, differ from their evolutionarily related sequences in terms of both amino acid sequence and structure. Thus, information from MSA would add noise in this regard.

Therefore, further improvements should be possible by not using MSA to predict the structure. As a result, prediction without MSA has reduced noise; however, it also means that there is no support from evolutionarily related sequences. It has been reported that AF2 retrained using a single sequence as input was inaccurate (Wu, et al., 2022). Thus, we used predicted structures which, although not perfect, were functional for input. Consequently, we trained our model to refine these predicted structures in a physically preferable situation. In selecting the training samples, we avoided segments that did not interact well with other segments and the training samples were penalized based on the number of residue-residue contacts. This procedure was based on the assumption that if a structure is not globular, if it lacked a sufficient internal interaction network, then the structure is flexible and machine learning methods cannot adequately model the rules of structures. In this regard, our model may be heavily biased toward globular structures but can still provide a useful baseline for predictions.

Here, we present a fine-tuned version of AlphaFold-Multimer, named AFM-Refine-G, which uses multimer structures of proteins as input and outputs refined structures. Roney and Ovchinnikov demonstrated that the official parameters of AlphaFold2 can be used to improve the structures without MSA features(Roney and Ovchinnikov, 2022), and our in-house benchmarks showed that fine-tuning the official parameters of AlphaFold-Multimer could be effective rather than training from scratch (data not shown). Therefore, we did not train the model from scratch but, instead, fine-tuned the official parameter param_multimer_model_1_v2. The performance of AFM-Refine-G was tested using two datasets, Ghani_et_al_Benchmark2 and Yin_et_al_Hard, which were adapted from studies by Ghani et al. (Ghani, et al., 2022) and Yin et al. (Yin, et al., 2022). The Ghani_et_al_Benchmark2 dataset consists of 17 recently published heteromers and the Yin_et_al_Hard dataset consists of 133 multimers, including immune-related complexes and repebody-antigen complexes, with many of complexes that AlphaFold-Multimer could not predict correctly. Firstly, the protein complexes in the test datasets were predicted using AlphaFold-Multimer’s official pipeline, and the predicted structures were fed into the AFM-Refine-G. Then, the input and refined structures were assessed in terms of DockQ and TM-score. For Ghani_et_al_Benchmark2, our model improved the quality of the predicted structures in general, whereas for Yin_et_al_Hard, some structures decreased in quality. In addition, 97 out of 133 structures in Yin_et_al_Hard were classified as “Incorrect” by CAPRI criteria, indicating substantial room for improvement in multimer predictions.

We further analyzed more recent targets, CASP15_multimer and Yin_and_Pierce_af23, using AlphaFold-Multimer version 2.3. With these two datasets, the tendency was similar to that of previous two datasets, suggesting that our approach (i.e., fine-tuning models with only sequence and precomputed structure as input) is still effective for new models. However, regarding the number of models assigned to the acceptable or higher CAPRI criteria, the difference were modest. Notably, in Yin_and_Pierce_af23, which consists of antibody-antigen and nanobody-antigen complexes, more than 50% of targets were classified as “Incorrect” by CAPRI criteria, highlighting the need for improvement in these challenging targets.

## Materials and Methods

### Training Script

The codes of AlphaFold-Multimer for the loss calculation and model architecture were copied from the inference code of DeepMind’s official GitHub repository downloaded on 2022-11-05. v2.2 code was cherry-picked and introduced. The codes were then modified to allow mixed precision. A bug for residues with the same residue index was resolved and “NotImplementedError” in structure module loss was commented out. Multi-chain permutation was considered when providing the ground truth. Other codes required for training, such as processing training samples or updating parameters, were written in-house. Scripts were executed as maximally deterministic (setting random seeds and environment variables, such as XLA_FLAGS=--xla_gpu_deterministic_reductions and TF_CUDNN_DETERMINISTIC=1). The chain center-of-mass loss term and clash loss term were not implemented.

### Contact-Based Spatial Cropping

To use physically preferred structures as the ground truth, we performed customized spatial cropping (Algorithm 1). We intended to obtain segments that 1) had a sufficient number of residues contacting (distance between two CA < 8.0 □) with other segments and 2) were longer than the predefined length. The algorithm could generate samples that did not meet these criteria and these were filtered after the multi-chain permutated ground-truth process (described below).

**Algorithm 1.**
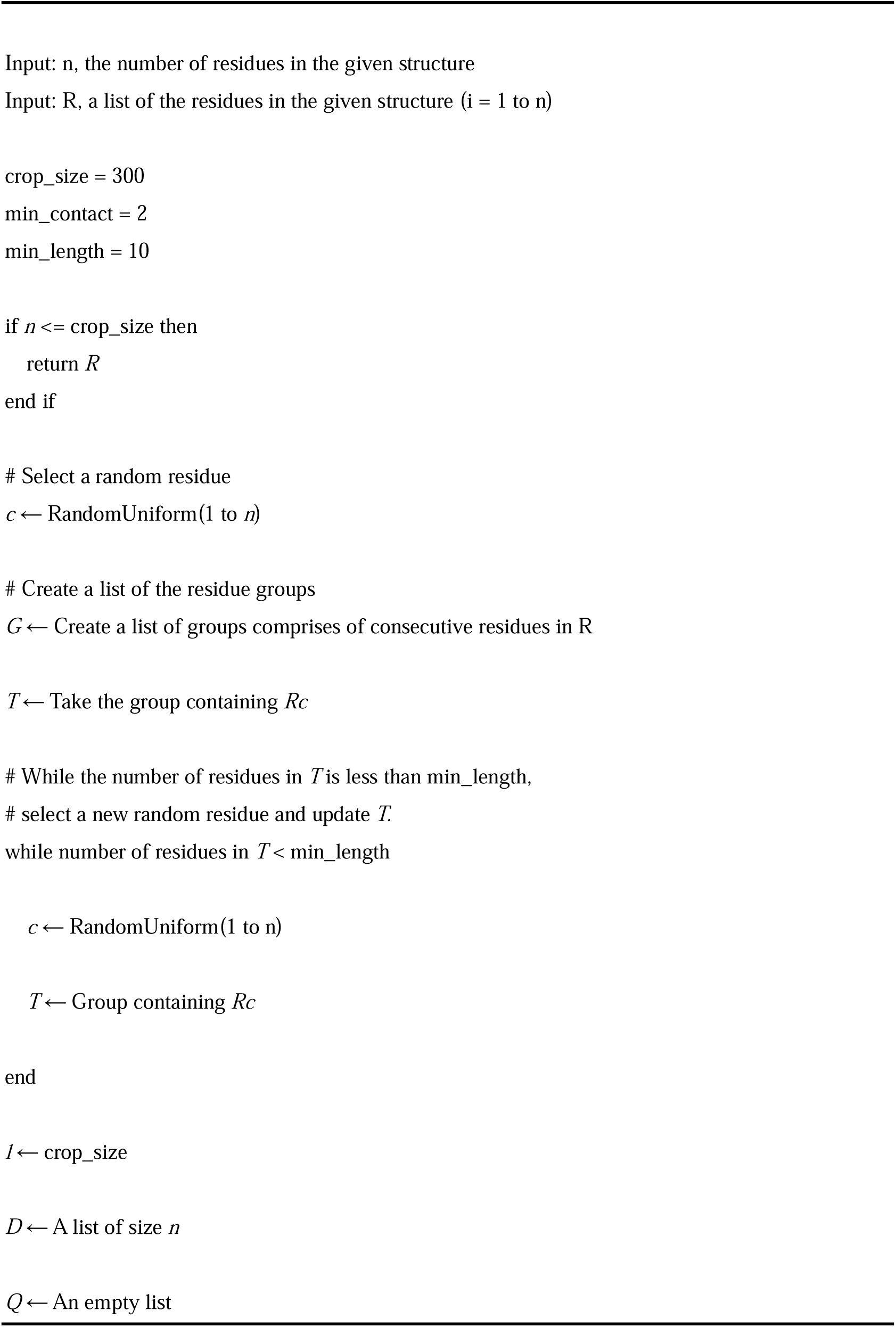
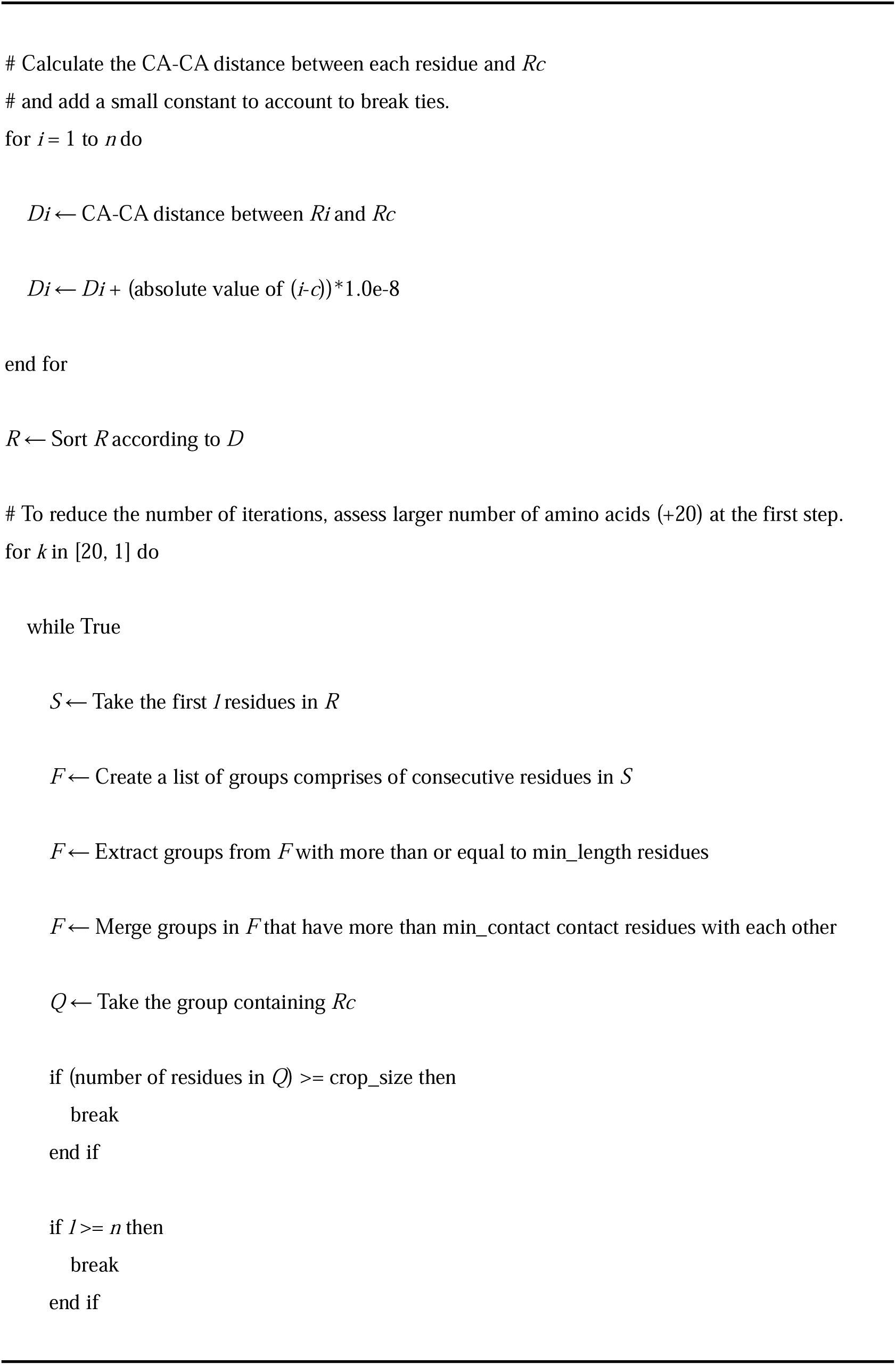
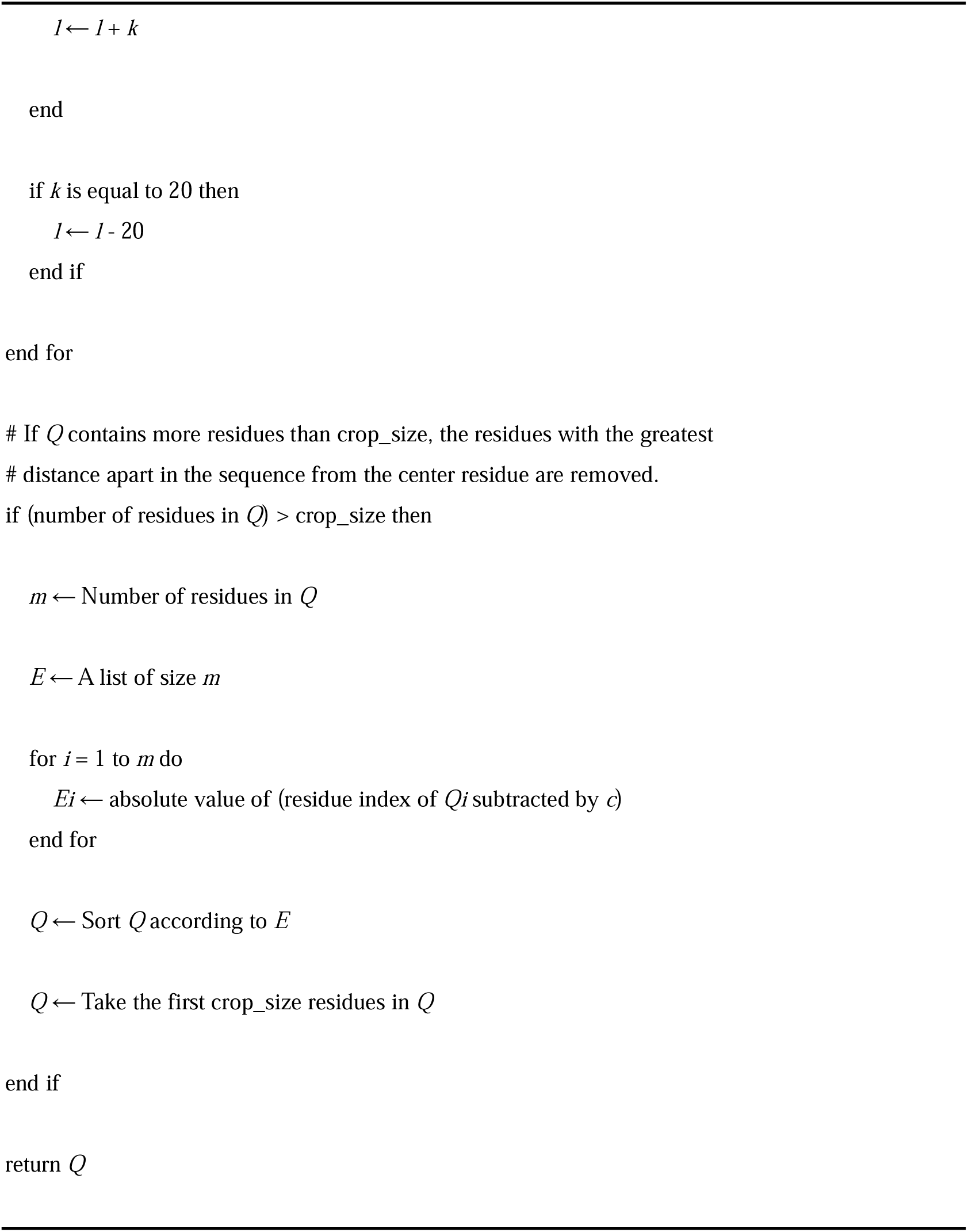
Contact-Based Spatial Cropping

### Multi-Chain Permutated Ground Truth

After spatial cropping, we generated ground-truth candidates by considering all multi-chain permutations. We filtered out ground truth candidates where any of their segments had residues contacting less than two other segments or consisting of less than 10 amino acids. At the training stage, predicted structures were superposed using the SVDSuperimposer from Biopython (Cock, et al., 2009), and a ground truth candidate with the smallest rms value was used for loss calculation.

### Non-Globular Penalty

In addition to the square root penalty for short sequences introduced in the original training procedure of AlphaFold2 (Jumper, et al., 2021), we introduced a penalty for nonglobular samples. This was intended to penalize entries that should be complexes but registered as monomers or incomplete complexes. These samples should have fewer contacts compared with entries that registered as complete complexes with the same number of amino acids.

If protein monomers or complexes are globular (spherical), the number of contacts becomes proportional to *N – N^2/3^*. Therefore, we calculated the normalized number of contacts as follows:

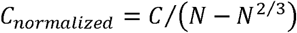

where *C* is the number of contacts, and *N* is the number of amino acids.

The plot of *C_normalized_* for the structures used in Epoch 1 is shown in Figure S1. The penalty for nonglobular samples was calculated as follows:

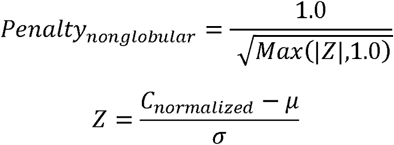

where *Z* is the Z-score, μ is the average of the normalized number of contacts, and σ is the standard deviation of the normalized number of contacts. μ and σ were calculated using the ground-truth structure candidates (after cropping) in Epoch 1, where μ = 5.713042053300396 and σ = 0.34169036005131537. The final loss for each sample was multiplied by both the short-sequence and non-globular penalties. We also penalized samples that had a large normalized number of contacts because they could be erroneous or have extremely rare properties.

### Training Dataset

The structural data in PDB (Berman, et al., 2000) were downloaded from the pdbj (Bekker, et al., 2022) ftp site (ftp.pdbj.org::ftp_data/biounit/PDB/divided/) on 2021-11-01. The .cif.gz files containing annotations were downloaded from ftp.pdbj.org::ftp_data/structures/divided/mmCIF/ on 2021-11-01. Entries that met the following criteria were included: 1) their release dates were before 2018-04-30, 2) resolutions were better than 9.0 □, 3) _entity_poly.type of the all polymer entities were polypeptide(L), 4) sequence lengths were less than or equal to 1024, 5) sequence lengths without ‘X’ or more than 4 consecutive ‘H’ were longer than 20, and 6) assembly data were provided in PDB format.

The chains were clustered with 40% sequence identity using the MMseqs2 (cloned from https://github.com/soedinglab/MMseqs2 on 2021-11-19) easy-cluster command (Steinegger and Soding, 2017). If the members in an entry belonged to different clusters, then those clusters were merged further. We discarded assembly entries with more than 2048 total amino acids or where the number of permutations was greater than 12. A total of 0.95 of clusters were used as the training dataset and 0.05 as the validation dataset. In each epoch, one entry was randomly selected from each cluster. Then the selected entries were processed with contact-based spatial cropping described above. If the cropped structures did not meet the criteria, they were discarded. Consequently, about 14,500 entries (samples) were used for training in each epoch.

### Input structures for training

We generated input structures using AlphaFold-Multimer and a customized protocol that reduced computation time and resources. MSA construction was performed using HHBLITS (v 3.3.0) (Steinegger, et al., 2019) against the uniclust30 (uniclust30_2018_08)(Mirdita, et al., 2017) database. The search was repeated up to three times using a query or the output, a3m, from the previous iteration. Iterations were stopped if the number of sequences in output a3m exceeded 200 after filtering (using hhfilter with options -cov 30 -id 90). The template database was downloaded on 2021-11-01 and the maximum template date was set to 2018-04-30. kalign (Lassmann, 2019) was not used because it may unexpectedly continue to run for a long time. Structures were predicted using five official multimer weights (v2), where they were selected randomly. The number of recycling steps was randomly selected between 0-3. If a sequence contained OX or TaxID attributes in its description line then these attributes were used for pairing.

### Training Procedure

The official weight param_multimer_model_1_v2 (CC-BY 4.0, Copyright 2021 DeepMind Technologies Limited, downloaded from https://storage.googleapis.com/alphafold/alphafold_params_2022-03-02.tar on 2022-03-11) was used as the source for the fine-tuned model. The atom coordinates of the predicted structure were inserted into the prev_pos of the input feature. Template embedding was not performed in this case. The single sequence data was fed to extra_msa and msa features. Mixed precision with float16 was used. Loss was scaled dynamically so that it became as large as possible but did not produce inf or nan in gradients. The Adam optimizer with warmup_exponential_decay_schedule of optax (Babuschkin, et al., 2020) was used for the parameter update. In the first step, the model was trained for one epoch. In the second step, the model was trained for 5 epochs. The other settings are listed in Table S1.

### Evaluation Procedure

#### Test Dataset

We used four benchmark sets to test the prediction performance of AFM-Refine-G. Ghani_et_al_Benchmark2 was collected from the benchmark 2 dataset introduced by

Ghani et al. (Ghani, et al., 2022), which consisted of heterodimeric complexes released after May 2018. Yin_et_al_Hard was collected from the Tables S4-S8 of the paper by Yin et al. (Yin, et al., 2022). The dataset consists of immune-related complexes and repebody-antigen complexes, that AlphaFold-Multimer version 2.2 mostly failed to predict. Entries released before 2018-4-30 were eliminated. The 6uk4 structure was not used because the interface between the antibody and antigen was not found in any biological assembly.

The CASP15_multimer dataset comprises 8 homomers or heteromers that were multimer targets in CASP15 experiments(Kryshtafovych, et al., 2023). The targets were included only if they met two criteria: 1) the total number of amino acids was less than 1024, and 2) the ground truth structures were available. The ground truth structures were downloaded from the CASP website or from the PDB if the structures were not available on the CASP website but PDB IDs were provided. Targets were excluded if their SEQRES record differed from the CASP query sequence. Note that T1119 is a repeat protein and has a large unmodelled region (PDB ID: 7SQ4), which may result in inaccurate scores.

The Yin_and_Pierce_af23 consists of 39 antibody-antigen or nanobody-antigen complexes extracted from the previous study by Yin and Pierce(Yin and Pierce, 2024), which were released after the cutoff of date of AlphaFold-Multimer version 2.3’s training date. Following Yin and Pierse’s approach, we used only variable domain for antibodies.

#### Predictions of Input Structures

For input structures of Ghani_et_al_Benchmark2 and Yin_et_al_Hard, we used the AlphaFold2 repository (https://github.com/google-deepmind/alphafold) cloned on 2022-09-09. For CASP15_multimer and Yin_and_Pierce_af23, we used more recent version of the AlphaFold2 repository cloned on 2024-08-25. The download dates for the required databases are listed in Table S2. For target 6tej, jackhmmer produced an extremely large number of hits, so we used only SwissProt instead of SwissProt+TrEMBL for MSA generation. For Ghani_et_al_Benchmark2 and Yin_et_al_Hard, we generated one structure per model without relaxation. For CASP15_multimer and Yin_and_Pierce_af23, we generated five structures per model. The predicted structures (5 or 25 per target) were used as inputs for AFM-Refine-G.

#### Collection of Experimental Structures

The structures were downloaded from the pdbj ftp site (ftp.pdbj.org::ftp_data/assemblies/mmCIF/divided) on 2022-09-09 for Ghani_et_al_Benchmark2 and Yin_et_al_Hard and 2024-08-23 for structures of CASP15_multimer (except for the structures which could be downloaded from CASP website) and Yin_and_Pierce_af23. When there were multiple entries (files), the file labeled “assembly1” was used. Chains were grouped according to their description, number of contacts with other chains, and visual inspections with PyMOL (Schrodinger, 2015) to find important interfaces (e.g., antibody-antigen interface). If there were multiple groups with the same types of interfaces, the groups that had receptor chains appearing earlier in the parsed structure by MMCIFParser of Biopython were used. We didn’t remove missing regions from query sequence.

#### Evaluation Measures

For Ghani_et_al_Benchmark2 and Yin_et_al_Hard datasets, we used DockQ (Basu and Wallner, 2016) and TM-score (Zhang and Skolnick, 2004) to evaluate the quality of the predicted models. DockQ provides a balanced assessment for multimer models, which is aimed at matching the CAPRI (Duan, et al., 2020) criteria. The ranges of DockQ < 0.23, 0.23 <= DockQ < 0.49, 0.49 <= DockQ < 0.80, and 0.80 <= DockQ correspond to Incorrect, Acceptable, Medium, and High in CAPRI criteria, respectively. As mentioned above, the list of chain groups used as “receptor(chain1)” and “ligand(chain2)” when supplied to DockQ is shown in Table S3. TM-score provides a global assessment of the predicted structures. The application is usually used for the assessment of monomers, but it can also be used for multimers with the “-ter 0” option. All cases of multi-chain permutations were considered, and the highest scores were used.

For CASP15_multimer, we used DockQ version 2 and US-align with the option “-TMscore 7 -ter 0 -het 1”. For Yin_and_Pierce_af23, we used DockQ version 1 and US-align with the option “-TMscore 7 -ter 0 -het 1”.

To compute LDDT (Mariani, et al., 2013) for evaluating the mean absolute error of pLDDT (Jumper, et al., 2021), we used the lddt tool from the OpenStructure package(Biasini, et al., 2013). All multi-chain permutations were considered, and the chain-mapping with the highest LDDT was selected. The chains were concatenated, and LDDT was calculated using only CA atoms.

## Results and Discussions

The multimer structures of two datasets, Ghani_et_al_Benchmark2 and Yin_et_al_Hard, were predicted using the official AlphaFold-Multimer version 2.2 (AFMv2.2) pipeline. The number of multimer predictions per model was set to 1. The predicted structures were then fed to AFM-Refine-G. The number of recycling steps for AFM-Refine-G was set to 10, and intermediate structures were produced. As a result, 750 structures were produced for the input, and 8250 structures were produced as the refined output. Improvements in scores (ΔDockQ and ΔTM-score) were calculated by subtracting the values of the refined structures from those of the input structures. ΔDockQ as a function of the number of recycling steps are shown in Figure 1. As a result, the improvement saturated at approximately 2-6, and we decided to analyze only the results obtained with three recycling steps hereafter.

**Figure 1.**
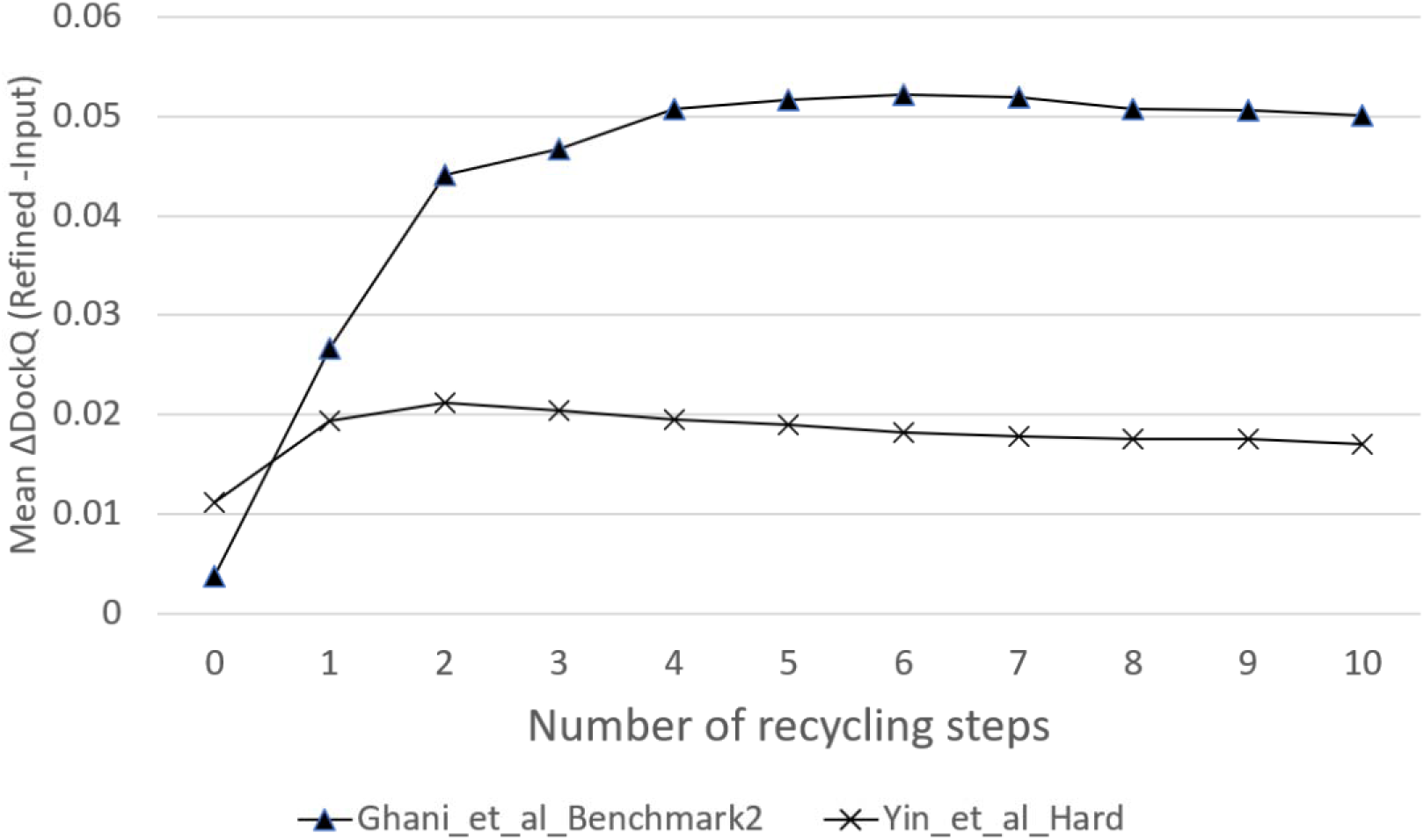
The mean DockQ as a function of the number of recycling steps. The ΔDockQ, DockQ of refined structures subtracted by that of input structures, calculated for all five structures per one target were averaged. A triangle indicates the result of Ghani_et_al_Benchmark2 (85 structures) and a cross indicates the result of Yin_et_al_Hard (665 structures).

### Improvement of Accuracy

Comparisons between the input and refined structural qualities are shown in Figure 2. Multiple models improved in quality with a ΔDockQ (Refined-Input) > 0.05 with 14 out of 85 and 101 out of 665 for Ghani_et_al_Benchmark2 and Yin_et_al_Hard, respectively. Furthermore, only a small number of models decreased in quality with ΔDockQ < −0.05, there were 0 and 14 for Ghani_et_al_Benchmark2 and Yin_et_al_Hard, respectively. The numbers of acceptable or higher quality predictions in CAPRI criteria were the same or increased (Table 1, Table S4). This indicates that the refinement was generally successful. However, there was a severe degradation of the TM-score of some samples in Yin_et_al_Hard (Figure 2d). The numbers of models with ΔTM-score > 0.05 were 5 and 18 for Ghani_et_al_Benchmark2 and Yin_et_al_Hard, respectively. Although no models had ΔTM-scores < −0.05 in Ghani_et_al_Benchmark2, 36 did in Yin_et_al_Hard. Of these 36 structures, 26 out of 36 structures were TCR-peptide-MHC complexes. We conducted a structural alignment of ground truth, input, and refined structures of 26 TCR-peptide-MHC cases using the TCR chains of 6r2l as templates (Figure S2). In the input structures, the global orientations of TCR and MHC chains were accurately predicted to some extent, but most of the positions of the peptides were not. In contrast, in the refined structures, positions of peptides were incorrect, too, and many of the MHC chains were also placed wrongly; as seen in 7n1f_model_5.refined, some of them showed aggregated forms. As illustrated in the ground truth structures in Figure S2, the global orientations of TCR and MHC are consistent across different entries, indicating that they can be learned during the training of models. While the TM-score measures global similarity, DockQ is a metric that also takes into account the prediction accuracy of interfaces that are of particular interest (e.g., the interface between the TCR and the peptide). The difference in degradation between DockQ and TM-score is likely due to differences in their scoring schemes. It is difficult to provide a clear answer to why AFM-Refine-G could not generate outputs that follow the global orientation. We should note that two technical factors may have contributed to this issue. Firstly, AFM-Refine-G employed a smaller crop_size (300) than the original AlphaFold-Multimer (384). Secondly, AFM-Refine-G was trained without incorporating chain centre-of-mass loss terms. However, it remains a matter of debate whether the global orientation of subunits should be maintained even if the prediction of interfaces that are of particular interest is inaccurate.

**Figure 2.**
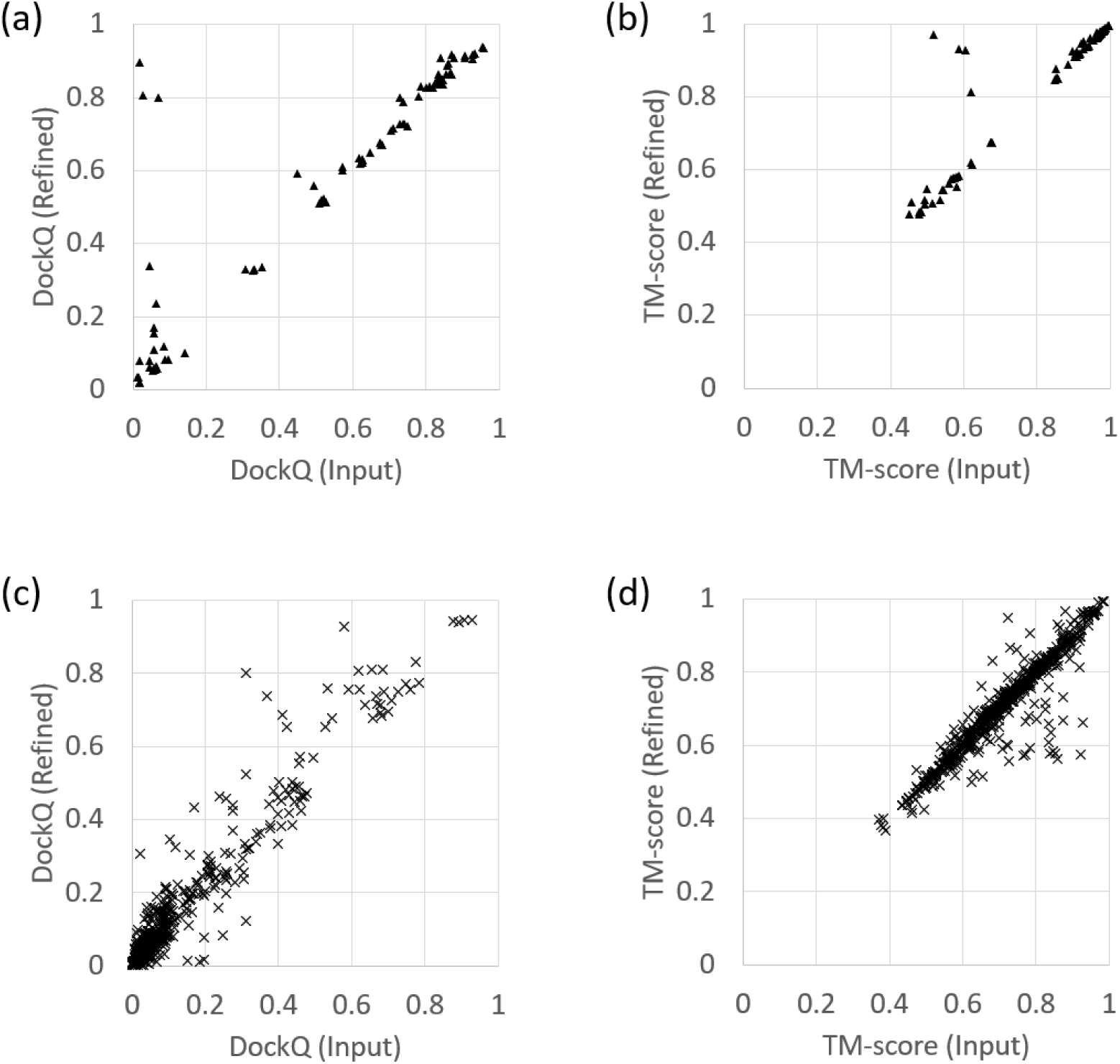
Comparison of structural quality between input and refined structures for Ghani_et_al_Benchmark2 and Yin_et_al_Hard datasets. (a) DockQ scores for Ghani_et_al_Benchmark2 dataset. (b) TM-scores for Ghani_et_al_Benchmark2 dataset. (c) DockQ scores for Yin_et_al_Hard dataset. (d) TM-scores for Yin_et_al_Hard dataset.

**Table 1.**
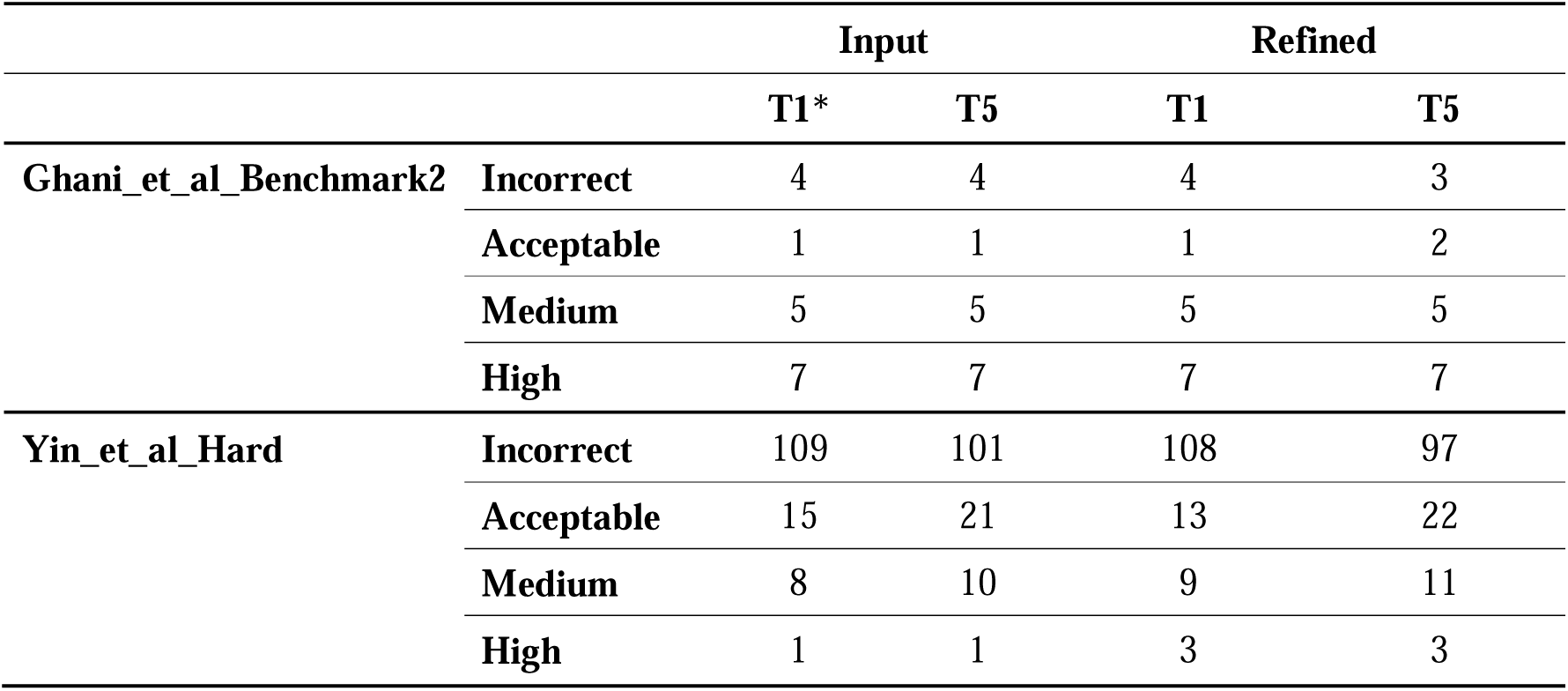
Number of targets assigned to CAPRI quality criteria. *T1 represents the top 1 structures based on the confidence metric (iptm*0.8+ ptm*0.2). T5 represents the structures with the highest DockQ scores among the all 5 structures.

### Predicted TM-score

AlphaFold-Multimer outputs structures and the confidence metric, weighted sum of interface predicted TM-score and predicted TM-score (iptm*0.8+ptm*0.2), to indicate the qualities of predicted structures and is used to rank them (Evans, et al., 2022). We analyzed whether this metric is still effective with AFM-Refine-G, specifically examining if structure quality estimation is possible without MSAs. The relationship between DockQ and (iptm*0.8+ptm*0.2) is shown in Figure 4. The R^2^ values between the metric and the DockQ or TM-score of the input and refined structures are listed in Table 2. The R^2^ values for refined structures were competitive, although slightly lower than that of input structures for Ghani_et_al_Benchmark2, slightly higher than that of the input structures for Yin_et_al_Hard. These results suggest that reasonable quality estimation is possible without MSAs when sufficient structural information is available.

**Figure 3.**
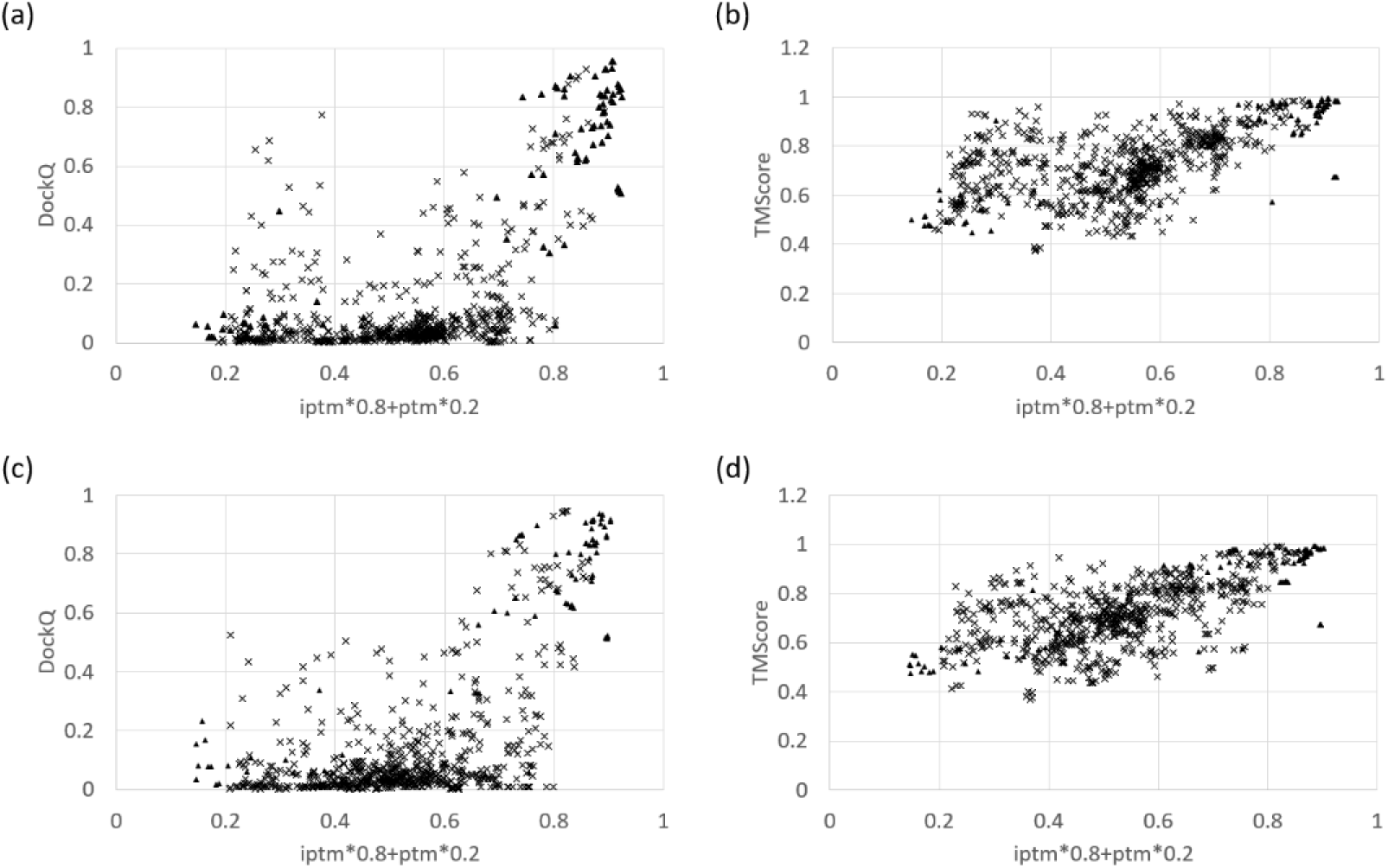
Correlation between the evaluation measures DockQ and TM-score and the confidence metric of AlphaFold-Multimer. Correlation between evaluation measures and confidence metric. The X-axis represents the weighted ipTM and pTM (iptm*0.8+ptm*0.2) utilized by AlphaFold-Multimer to rank its predicted models, while the Y-axis represents the DockQ or TM-score assessed using the predicted and experimental structures. Subfigure (a) shows the correlation between DockQ and iptm*0.8+ptm*0.2 of the input structures, while subfigure (b) shows the correlation between TM-score and iptm*0.8+ptm*0.2 of the input structures. Subfigure (c) displays the correlation between DockQ and iptm*0.8+ptm*0.2 of the refined structures, and subfigure (d) presents the correlation between TM-score and iptm*0.8+ptm*0.2 of the refined structures. The results for the Ghani_et_al_Benchmark2 dataset, comprising 85 structures, are indicated by triangles, while the results for the Yin_et_al_Hard dataset, comprising 665 structures, are represented by crosses. The R^2^ for each plot are provided in Table 2.

**Figure 4.**
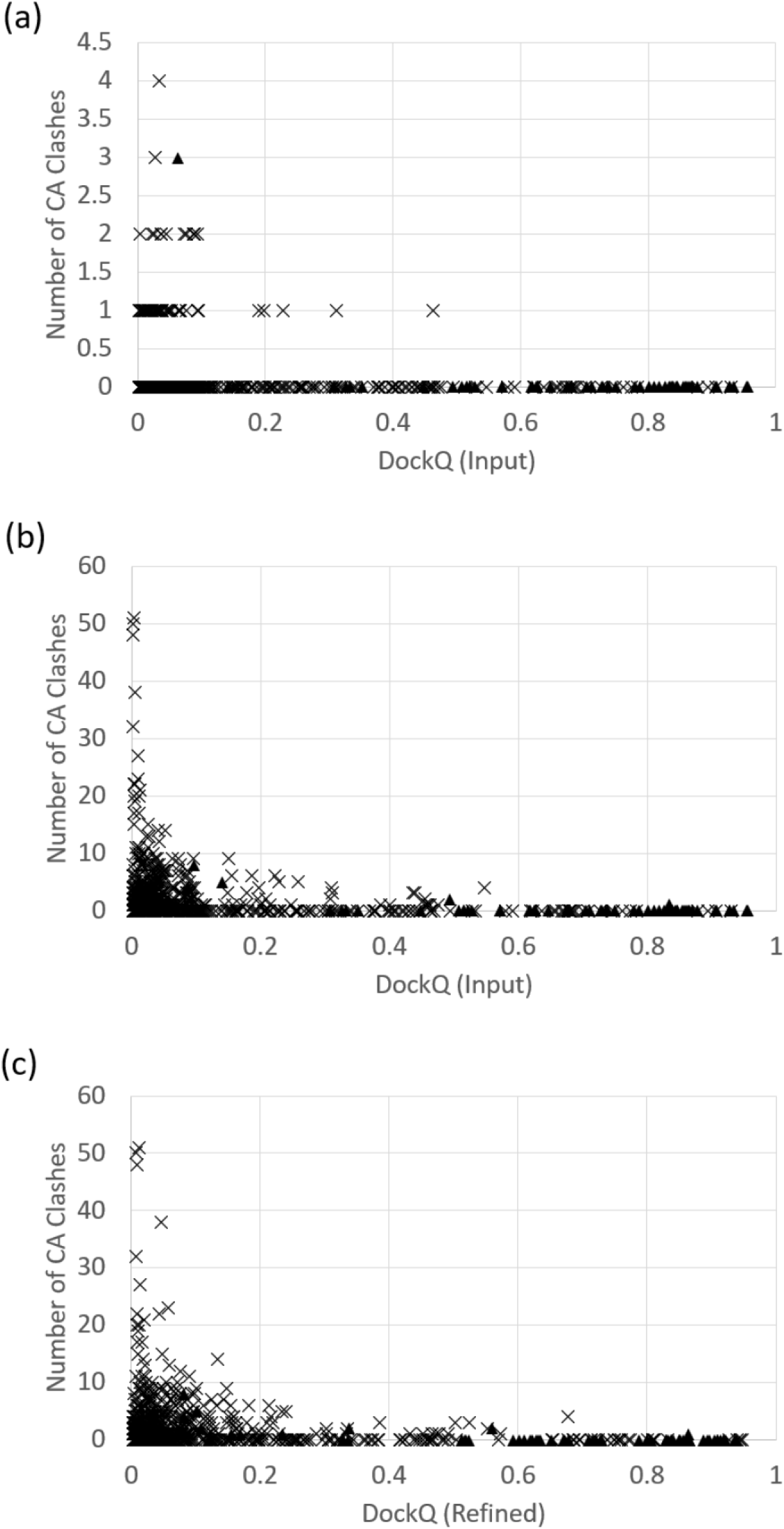
Number of CA atom clashes in the predicted models as a function of DockQ. A CA clash was defined as an instance where the distance between two CAs was less than 2.0 □. Subfigure (a) shows the number of clashes in the input structures as a function of the DockQ of the input structures. Subfigure (b) displays the number of clashes in the refined structures as a function of the DockQ of their corresponding input structures, and subfigure (c) presents the number of clashes in the refined structures as a function of the DockQ of the refined structures themselves.

**Table 2.**
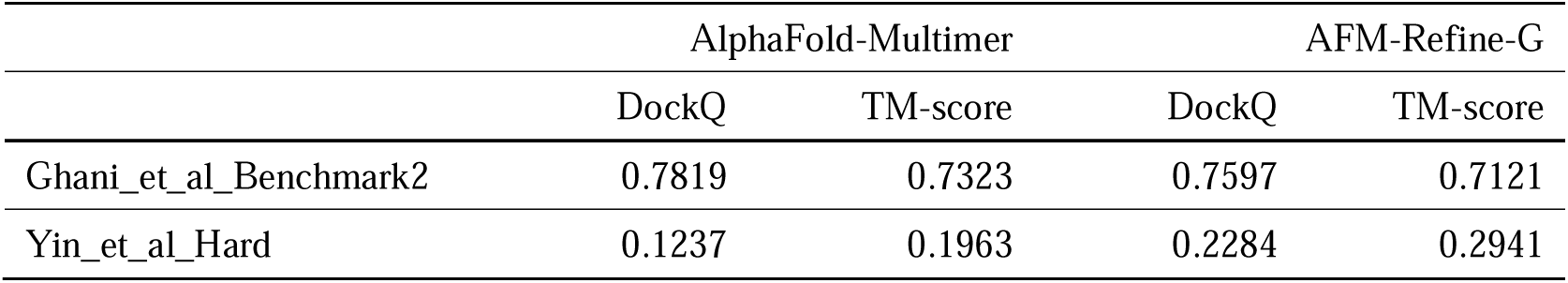
The R^2^ between the two evaluation measures and the confidence metric (iptm*0.8+ ptm*0.2) produced by AlphaFold-Multimer (version 2.2) and AFM-Refine-G.

|  | AlphaFold-Multimer |  | AFM-Refine-G |  |
| --- | --- | --- | --- | --- |
|  | DockQ | TM-score | DockQ | TM-score |
| Ghani_et_al_Benchmark2 | 0.7819 | 0.7323 | 0.7597 | 0.7121 |
| Yin_et_al_Hard | 0.1237 | 0.1963 | 0.2284 | 0.2941 |

### Atom Clashes

During the analysis, we found that there were severe atom clashes in the refined structures, which were rarely seen in the input structures. We counted the CA clashes (classified as when the distance between two CAs was less than 2.0 □) in all predicted structures. We then analyzed the relationships between CA clashes and the structure quality. The worst input structure had four CA clashes, and the worst refined structure had 51 CA clashes (Figure 4). The refined structures with a large number of clashes had a low DockQ in terms of refined and input structures (Figure 4B and C). 26 out of 750 structures had more than 10 CA clashes, and all had a DockQ less than 0.23, placing them in the ‘Incorrect’ prediction in CAPRI criteria. Several factors could cause this problem: 1) Intrinsic phenomena in our model. Because AFM-Refine-G was trained with samples biased toward globular proteins, the model might try to create globular structures if other good structures are not easily found. 2) Insufficient implementation of loss terms. We did not implement the updated loss terms which was not used in the first version of AlphaFold-Multimer, for example, the chain center-of-mass loss and clash loss terms (Evans, et al., 2022). 3) Differences in optimal loss weights. The weight for the structural violation loss of AlphaFold-Multimer (0.03) was much smaller than that of AlphaFold2 (1.0) (Evans, et al., 2022). It is possible that this value was not optimal for our model.

### Assessment Using More Recent Benchmark Set and an Updated AlphaFold Model

After the CASP15 experiment, DeepMind released a newer version of AlphaFold-Multimer, version 2.3 (AFMv2.3), which maintains the core network architecture but uses a more recent training dataset and larger crop size in training [https://github.com/google-deepmind/alphafold/blob/f251de6613cb478207c732bf9627b1e853c99c2f/docs/technical_note_v2.3.0.md accessed in 2024-12-26]. We investigated whether our refiner could improve models produced by this newer deep learning model. AFMv2.3 was run with the default pipeline settings without relaxation, and 25 predicted structures per target were fed to AFM-refine-G. As the test dataset, we used more recently published targets, named CASP15_multimer and Yin_and_Pierce_af23, 8 targets and 39 targets, respectively, results in 1175 predictions in total. The results are shown in Table 3, Table 4, Table S6, and Figure 5. The results were similar to those from previous benchmark sets with structures predicted by AFMv2.2. The accuracies of many structures were improved (Figure 5), indicating that our refiner performs consistently across model versions. However, some performance metrics showed degradation, such as the reduced number of acceptable predictions in CASP15_multimer T25 and medium quality predictions in Yin_and_Pierce_af23 T1 (Table 4), which may reflect the superior performance of the newly trained model.

**Figure 5.**
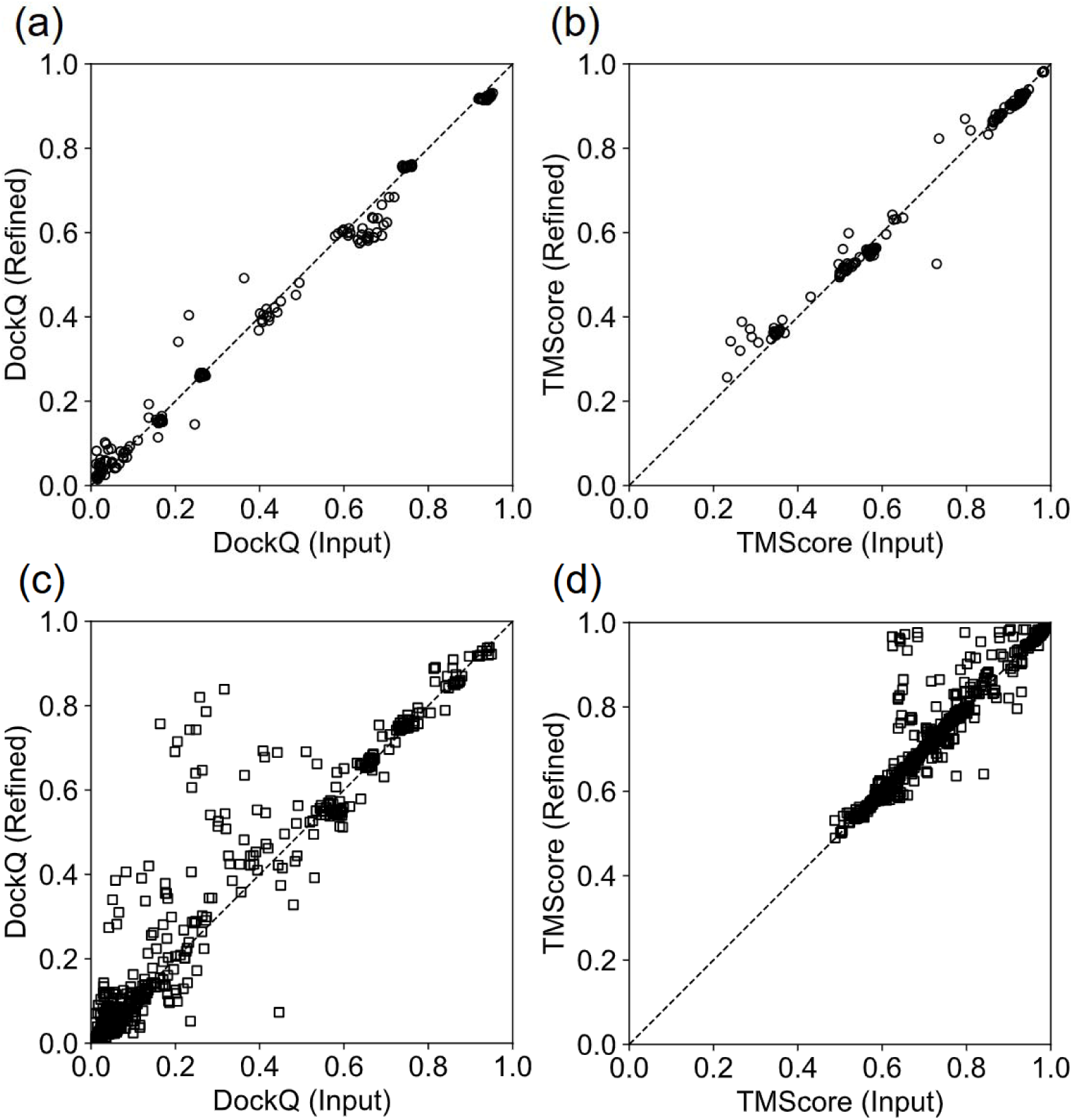
Comparison of structural quality between input and refined structures from recent datasets. Evaluation of all predicted models per target. (a) DockQ scores for the CASP15_multimer dataset. (b) TM-scores for the CASP15_multimer dataset. (c) DockQ scores for the Yin_and_Pierce_af23 dataset. (d) TM-scores for the Yin_and_Pierce_af23 dataset. CASP15_multimer structures are represented by circles and Yin_and_Pierce_af23 structures by squares.

**Table 3.** Number of structures with substantial changes after refinement for recent datasets. *Delta (Δ) values represent the difference between refined and input structure scores (refined - input)

|  | CASP15_multimer | Yin_and_Pierce_af23 |
| --- | --- | --- |
| $\Delta$ *DockQ > 0.05 | 7 | 81 |
| $\Delta$ DockQ < -0.05 | 14 | 26 |
| $\Delta$ TMScore > 0.05 | 9 | 41 |
| $\Delta$ TMScore < -0.05 | 1 | 13 |

**Table 4.** The number of targets assigned to CAPRI quality criteria for recent datasets. *T1 represents the top 1 structures based on the confidence metric (iptm*0.8+ ptm*0.2). T5 and T25 represent the structures with the highest DockQ scores among the top 5 and all 25 structures, respectively.

|  |  | Input |  |  | Refined |  |  |
| --- | --- | --- | --- | --- | --- | --- | --- |
|  |  | T1* | T5 | T25 | T1 | T5 | T25 |
| **CASP15_multimer** | **Incorrect** | 3 | 3 | 1 | 3 | 3 | 2 |
|  | **Acceptable** | 2 | 2 | 3 | 2 | 2 | 2 |
|  | **Medium** | 2 | 2 | 3 | 2 | 2 | 3 |
|  | **High** | 1 | 1 | 1 | 1 | 1 | 1 |
| **Yin_and_Pierce_af23** | **Incorrect** | 22 | 22 | 20 | 21 | 20 | 18 |
|  | **Acceptable** | 2 | 2 | 3 | 5 | 5 | 5 |
|  | **Medium** | 9 | 9 | 10 | 7 | 8 | 10 |
|  | **High** | 6 | 6 | 6 | 6 | 6 | 6 |

### Self-Accuracy Estimation

AlphaFold2 and AlphaFold-Multimer produce pLDDT scores that indicate the confidence of their per-residue predictions(Jumper, et al., 2021), which correspond to the predicted values of LDDT(Mariani, et al., 2013). Using the CASP15_multimer and Yin_and_Pierce_af23 datasets, we calculated the mean absolute error (MAE) of per-residue pLDDT for both the input and refined structures (Figure 6, Table S7). Some refined structures showed lower MAE, whereas others showed higher MAE. Overall, 657 out of 1175 cases had an increased MAE compared to their input structures, while the remaining cases showed a decrease. However, these changes were generally small and appeared to be target-specific. For example, of the 657 cases in which MAE increased, 20 predictions for just three targets increased MAE by more than 0.03, which may reflect a bias toward globular structures in our training procedure.

**Figure 6.**
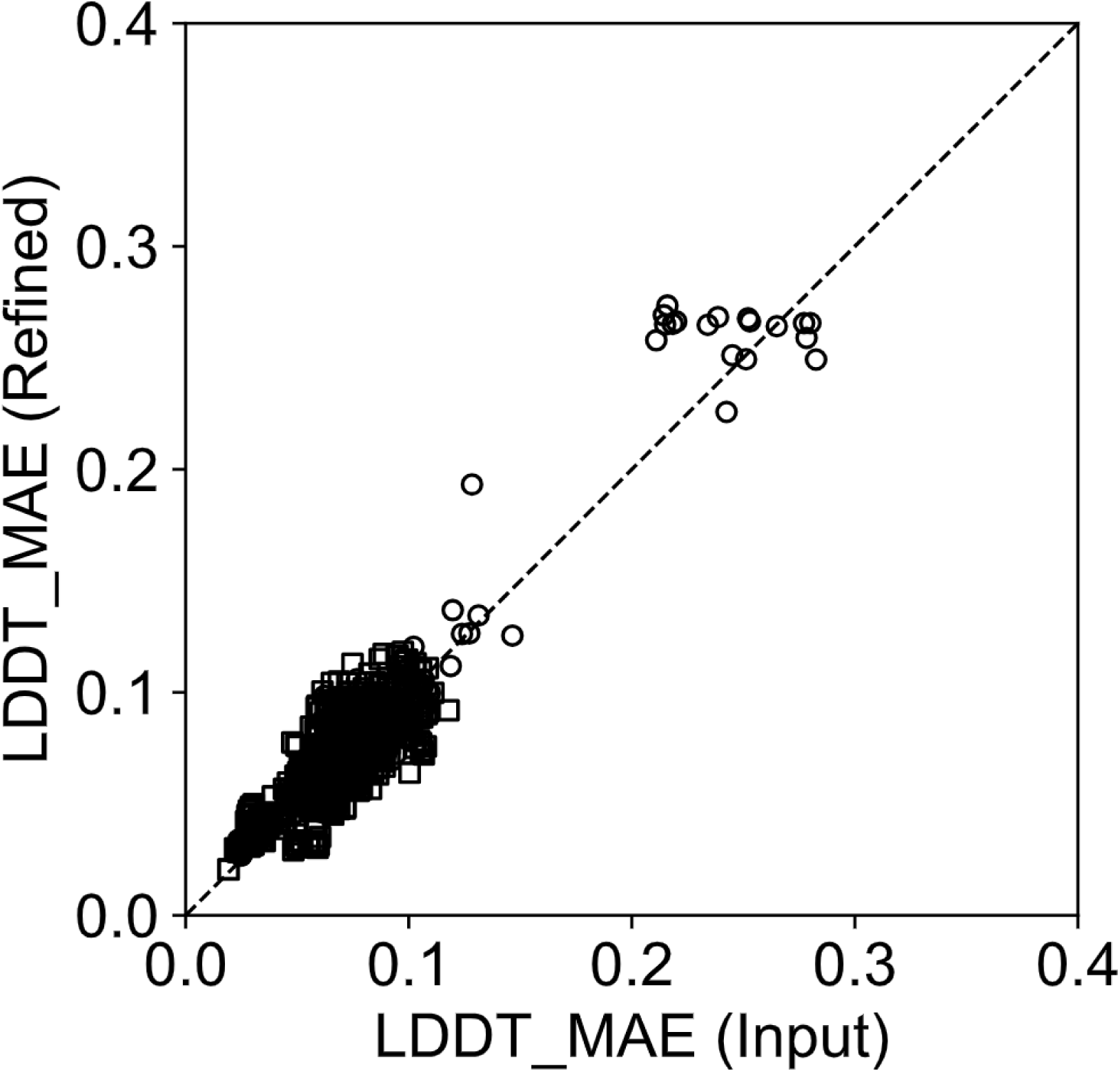
Performance of per-residue self-accuracy estimation. Comparison of mean absolute errors (MAE) between pLDDT and LDDT scores for input structures (x-axis) and refined structure (y-axis). Circle markers represent CASP15_multimer results and square markers represent Yin_and_Pierce_af23 results.

### Iterative Refinement

If the refinement model is effective, one might expect that feeding a refined structure back into the refiner would yield further improvements. To test this hypothesis, we used refined structures from CASP15_multimer and Yin_and_Pierce_2024 as input for additional refinement (Figure 7). We compared two approaches: performing refinement twice with 3 recycling steps each (hereafter referred to as “recycle 3×2”) versus performing refinement once with recycling steps 7 (hereafter referred to as “recycle 7”). Although some improvements were observed (Figures 7a and 7b), examining the datapoints around 0.8 on the x-axis in Figure 7c revealed that recycle 3×2 produced more markers compared to recycle 7 even for cases with DockQ < 0.23 (“Incorrect” predictions). Specifically, 20 predictions from recycle 3×2 and 1 prediction from recycle 7 had high confidence scores (iptm*0.8+ptm*0.2 > 0.8) despite low accuracy (DockQ < 0.23). This suggests that when an input structure matches the refinement model’s internal scoring function, it can generate misleadingly high confidence scores. Therefore, we recommend increasing the number of recycling steps in a single refinement process rather than performing iterative refinement.

**Figure 7.**
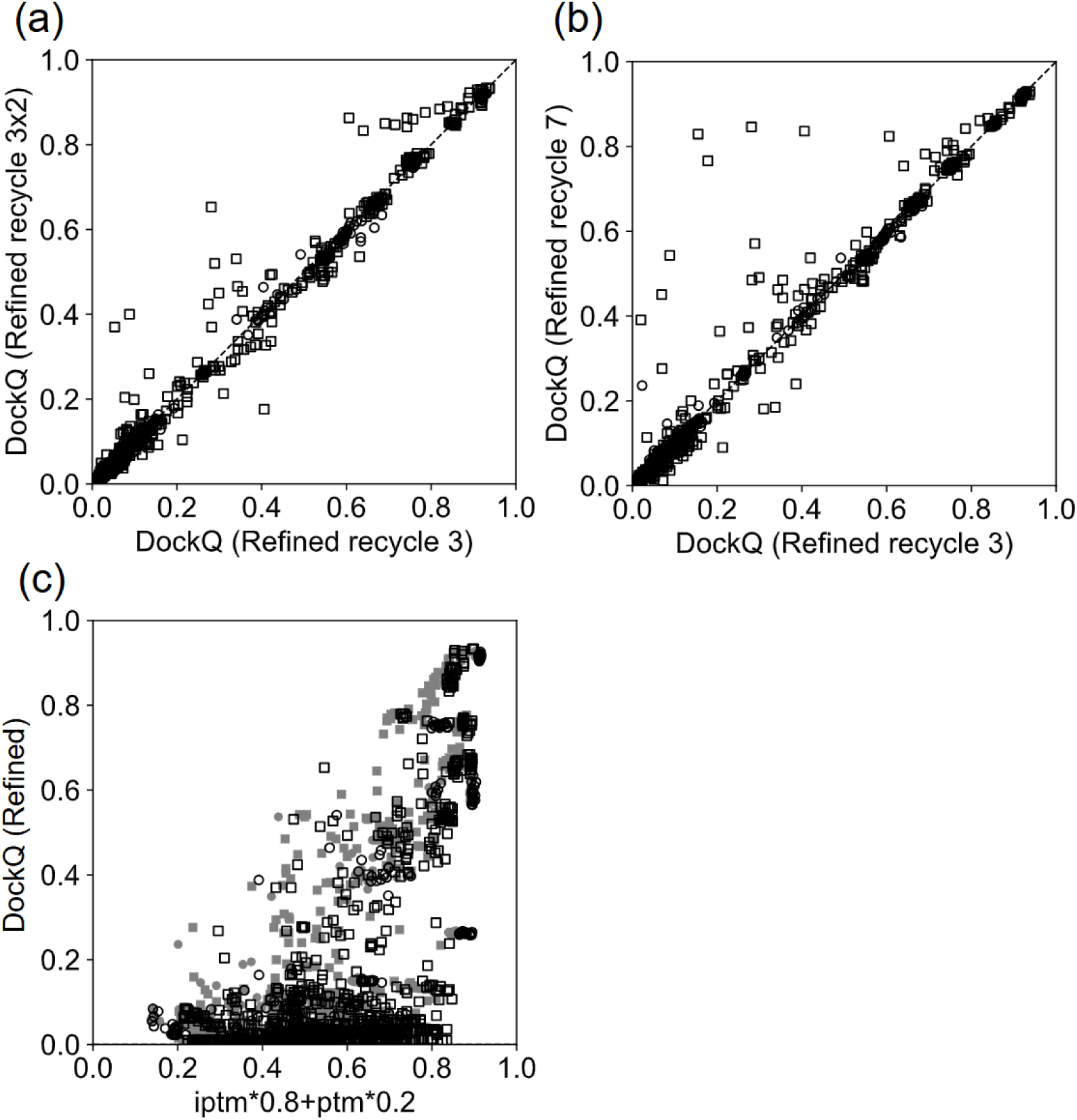
Effect of iterative refinement. Comparison of DockQ scores versus AlphaFold-Multimer confidence metric (iptm*0.8+ptm*0.2) for CASP15_multimer (circles) and Yin_and_Pierce_af23 (squares). (a) Comparison of DockQ scores between double refinement (two iterative refinement with 3 recycling steps each) and single refinement (one iterations with 3 recycling steps). (b) Comparison of DockQ scores between single refinement with 7 recycling steps and single refinement with 3 recycling steps. (c) Correlation between DockQ scores and AlphaFold-Multimer confidence metric. Gray filled markers represent results from single refinement with 7 recycling steps, and white markers with black outlines indicate results from double refinement with 3 recycling steps each.

## Conclusions

We hypothesized that AlphaFold-Multimer, which utilizes MSAs and templates as input, may be constrained by these inputs. Based on this hypothesis, we developed AFM-Refine-G, a fine-tuned version of AlphaFold-Multimer that refines multimeric structures without using MSAs and templates. While our refinement model improved predicted structures in many cases (DockQ improvement > 0.05 was found in 203 cases out of 1925 in total), the small differences in the number of highest-scoring models assigned to each CAPRI criterion (Tables 1 and 4) indicate that models of similar quality existed in the pools of unrefined structures. It suggests that the default AlphaFold-Multimer network had already learned a biophysical energy function beyond co-evolutionary information, as suggested by Roney and Ovchinnikov(Roney and Ovchinnikov, 2022). Furthermore, both the default AlphaFold-Multimer and our refinement model showed lower performance for immune-related targets, including antibody-antigen and nanobody-antigen complexes, compared to general targets, indicating room for improvement remains. While recent state-of-the-art structure prediction models (Abramson, et al., 2024) have shown improved performance for antibody-antigen complexes (though still less accurate than for general complexes), exploring these advanced approaches represents a promising direction for future work.

## Acknowledgements

We are grateful to Google DeepMind for developing AlphaFold2 and AlphaFold-Multimer, and to DeLano Scientific, LLC and Schrödinger, Inc. for providing and maintaining PyMOL. We also thank the many developers and maintainers for making various tools and databases available under generous licenses. Finally, we appreciate the assistance of ChatGPT, Claude 3.5, and DeepL in improving the English text.

## Conflict of Interest

The author declares no competing interests.

## Notes

### Competing Interest Statement

The authors have declared no competing interest.

### Summary of Updates

In the Results section, we added three new subsections: 'Assessment Using More Recent Benchmark Set and an Updated AlphaFold Model', 'Self-Accuracy Estimation', and 'Iterative Refinement'. The corresponding part of the 'Evaluation Procedure' in the 'Materials and Methods' section was also updated. We revised the 'Supplemental Tables', 'Abstract', 'Introduction', and 'Conclusion' sections to reflect these new findings. In addition, minor wordings were changed, errors and typos were corrected. And the 'Acknowledgement' section was updated.

