## Supplemental Figures for "Refinement of AlphaFold-Multimer structures with single sequence input"

| Number of amino acids | Normalized number of contacts |
| --- | --- |
| 300 | 6.556 |
| 300 | 6.43452 |
| 300 | 6.556 |
| 300 | 6.43452 |
| 300 | 6.556 |
| 300 | 6.43452 |
| 300 | 6.556 |
| 300 | 6.43452 |
| 300 | 5.50187 |
| 300 | 5.39607 |
| 300 | 5.36472 |
| 300 | 5.33728 |
| 300 | 5.29026 |
| 300 | 5.2158 |
| 259 | 6.12269 |
| 300 | 5.8859 |
| 291 | 5.67412 |
| 290 | 5.66235 |
| 300 | 6.15238 |
| 300 | 5.44309 |
| 300 | 5.81537 |
| 300 | 5.81537 |
| 291 | 5.37868 |
| 300 | 5.81537 |
| 300 | 5.81537 |
| 143 | 5.16194 |
| 300 | 5.90158 |
| 234 | 5.67269 |
| 208 | 5.89955 |
| 300 | 6.04657 |
| 234 | 6.16752 |
| 300 | 5.91725 |
| 300 | 5.85064 |
| 300 | 5.85064 |
| 300 | 5.91725 |
| 300 | 5.91725 |
| 300 | 5.85064 |
| 248 | 5.34224 |
| 153 | 5.39416 |
| 300 | 5.41958 |
| 277 | 5.37724 |
| 291 | 5.48795 |
| 288 | 5.47488 |
| 300 | 5.97995 |
| 289 | 6.47402 |
| 300 | 5.73307 |
| 300 | 5.73307 |
| 156 | 5.44792 |
| 296 | 5.64024 |
| 300 | 5.79969 |
| 121 | 6.15313 |
| 121 | 6.15313 |
| 84 | 5.63099 |
| 84 | 5.63099 |
| 300 | 5.60376 |
| 300 | 5.58416 |
| 300 | 5.77618 |
| 293 | 5.36391 |
| 293 | 5.36391 |
| 300 | 5.49011 |
| 266 | 5.38196 |
| 160 | 5.63865 |
| 300 | 6.41885 |
| 300 | 6.41885 |
| 300 | 5.57241 |
| 300 | 5.90158 |
| 206 | 6.15943 |
| 300 | 5.88199 |
| 300 | 5.72132 |
| 155 | 5.39855 |
| 91 | 5.4544 |
| 300 | 5.4666 |
| 296 | 5.46137 |
| 300 | 5.9486 |
| 300 | 5.95644 |
| 246 | 5.73669 |
| 254 | 5.90015 |
| 135 | 5.77823 |
| 139 | 5.14414 |
| 300 | 5.70564 |
| 105 | 5.54727 |
| 300 | 5.99563 |
| 74 | 5.00231 |
| 300 | 5.22756 |
| 300 | 5.39607 |
| 300 | 5.39607 |
| 205 | 6.03291 |
| 271 | 5.38576 |
| 271 | 5.38576 |
| 120 | 5.63388 |
| 97 | 5.41582 |
| 87 | 5.40333 |
| 300 | 5.28242 |
| 300 | 5.2785 |
| 146 | 5.72404 |
| 300 | 5.60768 |
| 300 | 5.64686 |
| 235 | 5.88568 |
| 300 | 5.68997 |
| 300 | 5.73699 |
| 134 | 5.34254 |
| 144 | 5.86131 |
| 300 | 5.83888 |
| 298 | 5.85669 |
| 300 | 5.74091 |
| 300 | 5.74483 |
| 300 | 6.25818 |
| 300 | 5.83104 |
| 300 | 5.46268 |
| 300 | 5.47052 |
| 300 | 5.35296 |
| 162 | 5.69236 |
| 162 | 5.69236 |
| 162 | 5.69236 |
| 162 | 5.69236 |
| 162 | 5.69236 |
| 162 | 5.69236 |
| 297 | 5.13693 |
| 295 | 5.14988 |
| 295 | 5.14589 |
| 293 | 5.15096 |
| 293 | 5.15899 |
| 293 | 5.15096 |
| 111 | 5.41504 |
| 300 | 5.36863 |
| 300 | 5.86239 |
| 300 | 5.67821 |
| 300 | 5.65862 |
| 260 | 5.75563 |
| 260 | 5.75563 |
| 125 | 5.63 |
| 300 | 5.86239 |
| 298 | 5.85275 |
| 300 | 5.88199 |
| 300 | 5.68997 |
| 300 | 5.70956 |
| 300 | 5.87415 |
| 300 | 5.88982 |
| 300 | 5.70564 |
| 200 | 5.63931 |
| 200 | 5.63931 |
| 300 | 5.6547 |
| 210 | 5.79951 |
| 210 | 5.79951 |
| 300 | 5.25891 |
| 300 | 6.02306 |
| 150 | 5.67468 |
| 300 | 5.79969 |
| 300 | 5.63903 |
| 300 | 5.63903 |
| 300 | 5.79969 |
| 300 | 5.79969 |
| 300 | 5.63903 |
| 67 | 5.46493 |
| 300 | 5.85455 |
| 300 | 5.70956 |
| 292 | 5.72612 |
| 101 | 5.8629 |
| 101 | 5.8629 |
| 300 | 6.0309 |
| 300 | 6.05049 |
| 127 | 5.36694 |
| 86 | 5.80307 |
| 86 | 5.80307 |
| 125 | 5.72 |
| 300 | 5.25891 |
| 300 | 5.26675 |
| 103 | 6.05972 |
| 96 | 5.31758 |
| 300 | 6.14062 |
| 171 | 6.07738 |
| 300 | 5.32161 |
| 272 | 5.95601 |
| 300 | 5.8232 |
| 300 | 5.81929 |
| 263 | 6.16353 |
| 198 | 5.2125 |
| 198 | 5.2125 |
| 300 | 5.97603 |
| 296 | 6.03772 |
| 80 | 5.82743 |
| 300 | 5.82712 |
| 138 | 5.66061 |
| 138 | 5.66061 |
| 194 | 6.13753 |
| 300 | 5.60376 |
| 297 | 5.58449 |
| 288 | 5.79813 |
| 288 | 5.79813 |
| 300 | 5.20797 |
| 300 | 5.18054 |
| 267 | 6.1232 |
| 300 | 5.29026 |
| 300 | 5.90942 |
| 198 | 5.47465 |
| 198 | 5.47465 |
| 203 | 5.41971 |
| 300 | 5.68213 |
| 300 | 5.68213 |
| 204 | 5.54486 |

**Figure S1. A scatter plot of the normalized number of contacts. A dot represents one ground truth structure candidate after cropping.** The outlier dot at (81, 9.4734) is 3boi, entitled "Snow Flea Antifreeze Protein Racemate".

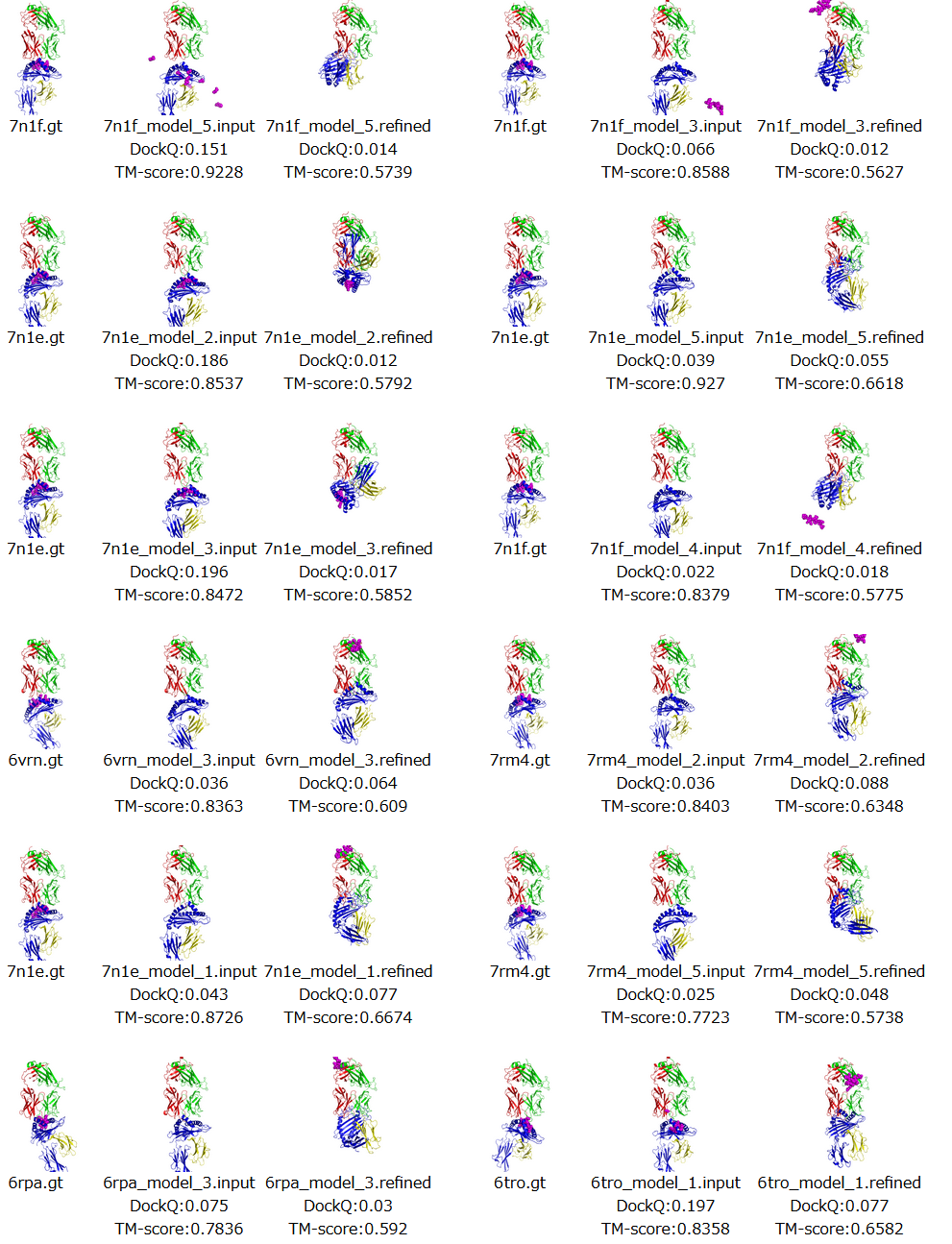

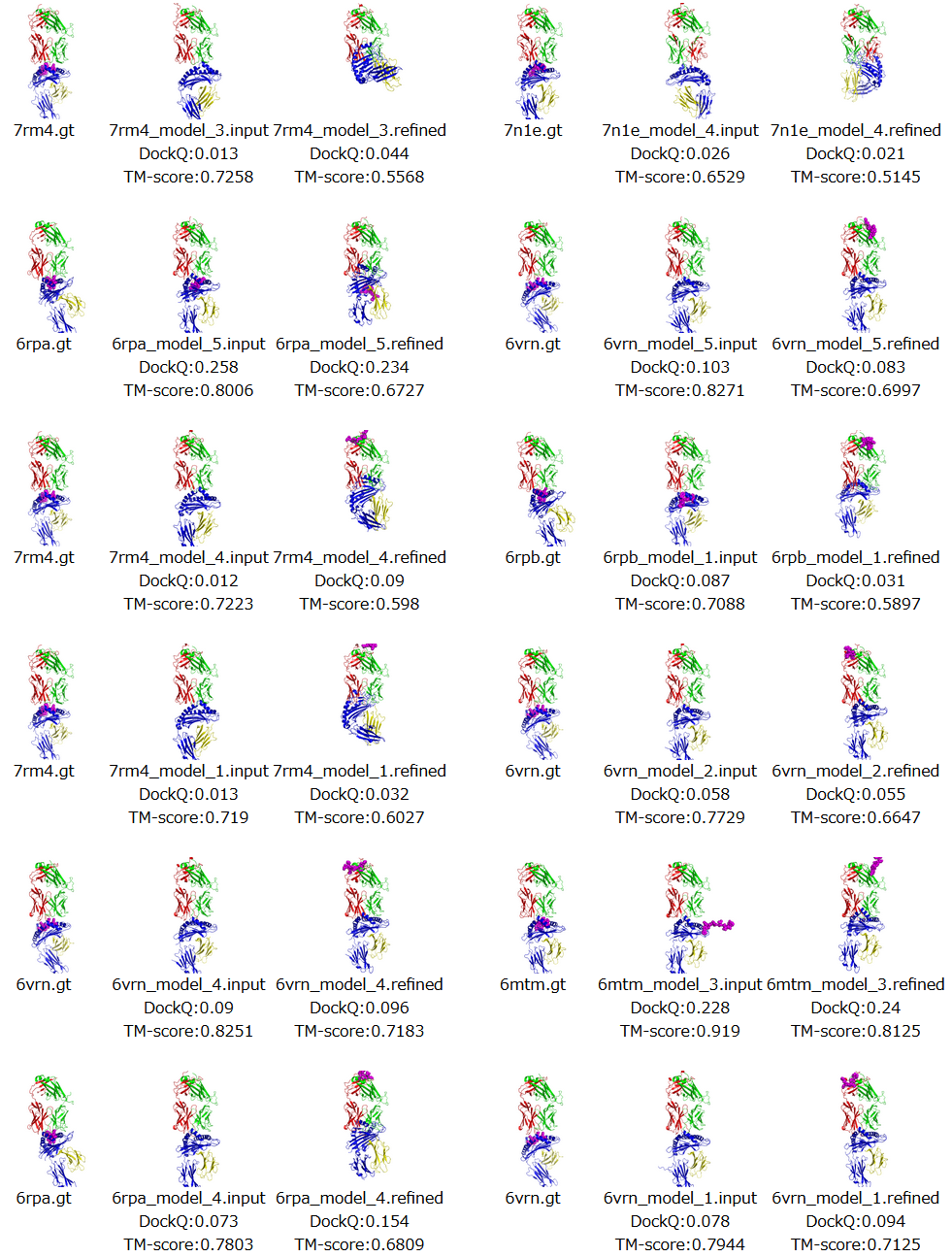

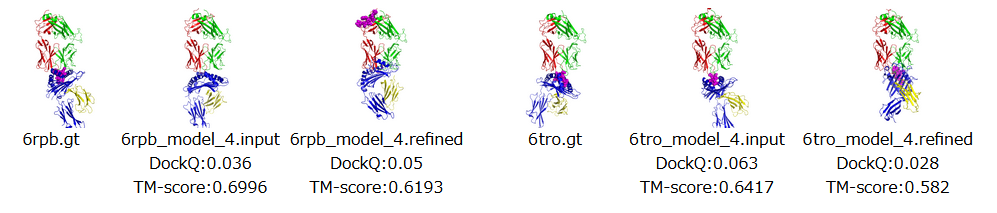

**Figure S2. The 26 TCR-peptide-MHC structures which had severe TM-score degradation.** The structures were sorted in ascending order of ΔTM-score and placed from the top left to the bottom right of the image. The structure alignment and image generation were done with PyMOL (Schrodinger, 2015). “.gt”, “.input”, and “.refined” indicates ground truth, input, and refined structures, respectively. model_X represents the model parameter used to produce the input structures. The peptides were drawn in magenta spheres. Some of the predicted peptides were out of the frame and are not visible in the image.
